# Determinants of the population health distribution, or why are risk factor-body mass index associations larger at the upper end of the BMI distribution?

**DOI:** 10.1101/537829

**Authors:** David Bann, Emla Fitzsimons, Will Johnson

**Author notes:** Contributed equally (Emla Fitzsimons, Will Johnson).

## Abstract

Most epidemiological studies examine how risk factors relate to average difference in outcomes (linear regression) or odds a binary outcome (logistic regression); they do not explicitly examine whether risk factors are associated differentially across the distribution of the health outcome investigated. This paper documents a phenomenon found repeatedly in the minority of epidemiological studies which do this (via quantile regression) -associations between a range of established risk factors and body mass index (BMI) are progressively stronger in the upper ends of the BMI distribution. In this paper, we document this finding and provide illustrative evidence of it in a single dataset (the 1958 British birth cohort study). Associations of low childhood socioeconomic position, high maternal weight, low childhood general cognition and adult physical inactivity with higher BMI are larger at the upper end of the BMI distribution, on both absolute and relative scales. For example, effect estimates for socioeconomic position and childhood cognition were around three times larger at the 90^th^ compared with 10^th^ quantile, while effect estimates for physical inactivity were increasingly larger from the 50^th^-90^th^ quantiles, yet null at lower quantiles. We provide potential explanations for these findings and discuss possible research and policy implications. We conclude by stating that tools such as quantile regression may be useful to better understand how risk factors relate to the distribution of health -particularly so in obesity research given conventional reliance on cut-points -yet for other outcomes in addition given the continuous nature of population health.

---

Epidemiology is concerned with understanding the distribution of health in a given population—first in describing it, and second in understanding its determinants.^1 2^ Yet in the majority of etiological applications, distributions are seldom of explicit focus regardless of the analytical tool used. Most papers investigating the determinants of body mass index (BMI) use either linear regression—to examine mean differences in BMI in different risk factor groups—or logistic regression—to examine if risk factor groups have higher odds of obesity. Neither of these options can straightforwardly whether risk factors are associated with differences across the distribution of the outcome in question (see Figure 1). Such differences may be anticipated—since the population BMI distribution has become increasingly right-skewed from the 1980s^3 4^ risk factors which have contributed to this may have had a disproportionately stronger effect at the upper end of the BMI distribution (and/or simply increased in prevalence).

**Figure 1.**
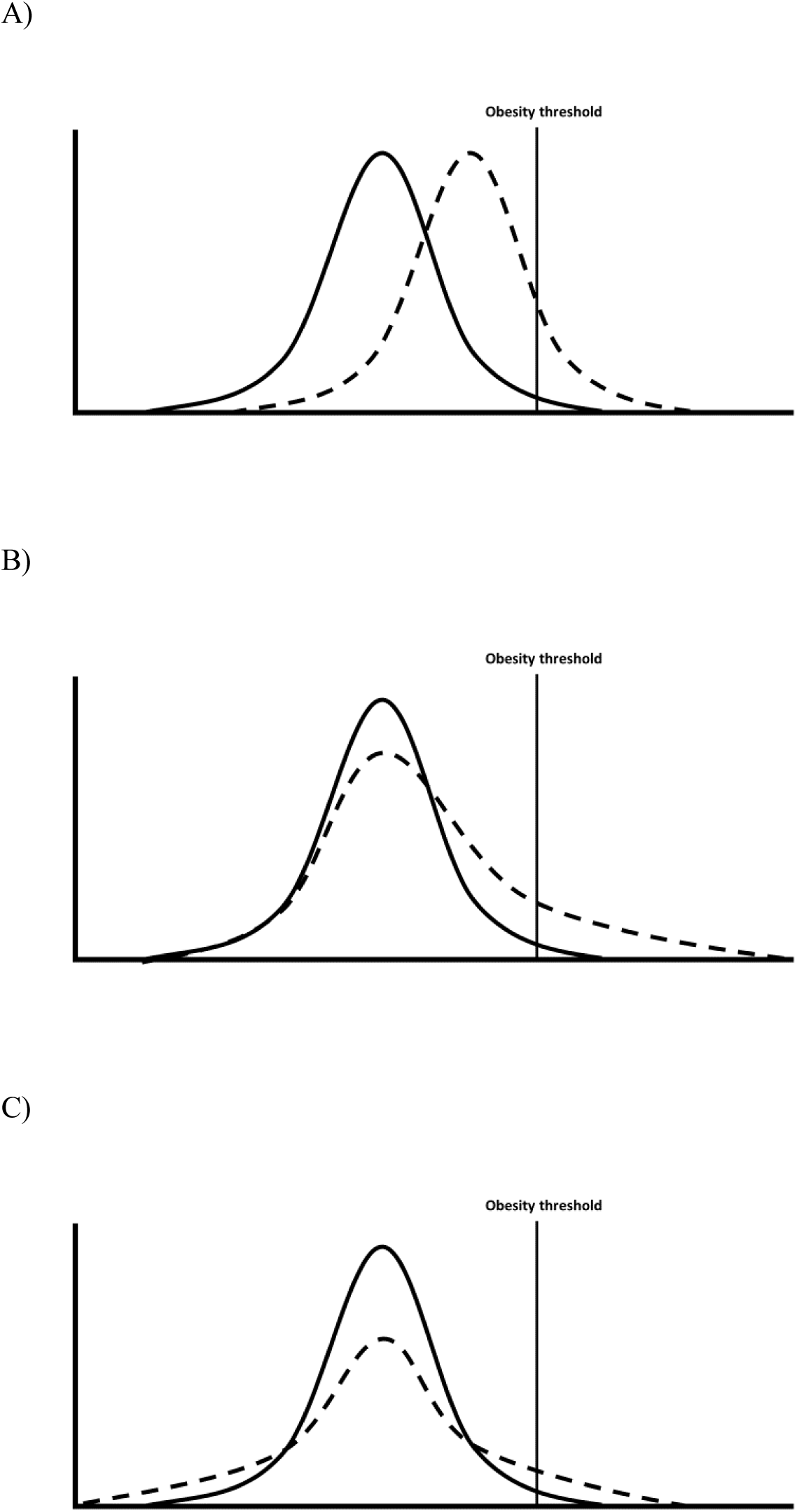
Comparisons between groups—mean differences only (A), mean differences driven particularly by differences at the upper quantiles (B); no mean difference yet different distributions (C), adapted from^5^

As noted by authors recently in the epidemiological literature,^5^ quantile regression is an one analytical tool which enables investigation of risk factor-outcome associations across the outcome distribution (ie, beyond standard cut-points). The statistical underpinning has been described previously elsewhere, as have applied examples of its interpretation, and (beyond the scope of the current paper) technical work on heterogenous treatment effects.^6-9^ Briefly, while linear regression estimates mean differences in outcomes across risk factor groups (which are likely identical in Fig 1A and B), and logistic regression compares odds of being above a threshold (odds ratios are both >1 in Fig 1A and B), quantile regression estimates the difference in a given quantile of the outcome distribution. For example, when comparing Fig 1B with Fig 1A, the median (50^th^ quantile) differences are likely similar, yet differences in the 90^th^ BMI percentile are notably higher Fig 1B.

Using quantile regression, we recently observed that absolute socioeconomic inequalities in children’s BMI were substantially larger in higher BMI quantiles;^10^ mean BMI differences in the lowest vs highest socioeconomic position (SEP) (in the cohort born in 2001 at 11y) was 1.3kg/m^2^ (95% CI: 0.9, 1.6); the median difference was 0.98 kg/m^2^ (95% CI: 0.63, 1.33), yet the difference at the 90^th^ percentile was 2.54 (1.85, 3.22). Similar finding have been observed in other studies in the UK, Spain, and Norway.^11-13^ Across the literature, there appears to be evidence for a phenomenon which does not seem to have been explicitly noted nor explained—in the minority of cases where the outcome distribution is explicitly investigated, associations between a myriad of risk factors and BMI are progressively larger at the upper ends of the BMI distribution. This includes genetic factors,^14 15^ behavioral factors (physical activity, sedentary behavior, and diet^16 17^), and family factors (maternal BMI or exercise).^16 18^ In this paper, we provide an illustrative example of this in a single dataset, provide potential explanations (substantive and methodological), and discuss potential implications for epidemiological research and policy.

## Demonstration

Data are from the 1958 British Birth cohort study, a longitudinal study described in detail elsewhere,^19^ with prospective risk factors data and BMI measured at 45 years. Table 1 shows associations between multiple established risk factors for high BMI: low childhood SEP (birth), high maternal weight (birth), low childhood general cognition (11y), and adult physical inactivity (42y). For each risk factor, the magnitude of associations was substantially larger at higher BMI quantiles. Associations of low SEP low cognition with higher BMI were around three times larger at the 90^th^ compared with 10^th^ quantile; effect estimates for maternal education were almost twice as large, while effect estimates for physical inactivity were null at lower quantiles and only found at higher BMI quantiles. Similar findings were found with waist circumference as an outcome, suggesting that findings are not an artifact caused by the indirect adiposity measure used (Appendix Table 1).

**Table 1.** Associations between risk factors for body mass index (kg/m^2^) at 45y using both linear regression and quantile regression in the 1958 British birth cohort study

| VARIABLES | Linear regression estimates<br>(mean difference in BMI) | Quantile regression estimates<br>(difference in BMI at below quantiles) |  |  |  |  |
| --- | --- | --- | --- | --- | --- | --- |
|  |  | q10 | q25 | q50 | q75 | q90 |
| Paternal social class (per 1 category lower), birth | 0.33***<br>(0.049) | 0.20***<br>(0.067) | 0.20***<br>(0.072) | 0.29***<br>(0.060) | 0.37***<br>(0.079) | 0.62***<br>(0.15) |
| Maternal weight (per 1 higher category), birth | 0.55***<br>(0.039) | 0.39***<br>(0.046) | 0.46***<br>(0.033) | 0.50***<br>(0.040) | 0.66***<br>(0.054) | 0.76***<br>(0.12) |
| General cognition (per 1 lower SDS), 11y | 0.41***<br>(0.064) | 0.18**<br>(0.087) | 0.32***<br>(0.081) | 0.41***<br>(0.088) | 0.51***<br>(0.078) | 0.67***<br>(0.15) |
| Physical exercise (inactive vs active), 42y | 0.66***<br>(0.13) | -0.22<br>(0.19) | 0.081<br>(0.14) | 0.59***<br>(0.21) | 1.11***<br>(0.15) | 1.61***<br>(0.38) |
| Observations | 6,943 | 6,943 | 6,943 | 6,943 | 6,943 | 6,943 |

Standard errors in parentheses
\*\*\* p<0.01, \*\* p<0.05, \* p<0.1
Note: models are mutually adjusted.

## Why could risk factor-outcome associations be stronger at the upper end of the distribution?

### 1. Heterogenous (non-constant) causal effects of a single risk factor

Risk factors may have multiple contrasting causal effects on the outcome which differ across the outcome distribution. For example, low SEP has been shown to be associated with increased risk of both obesity (prevalence ∼20%) and risk of thinness (prevalence ∼ 6%)^20^ such that quantile regression estimates might show a negative relationship with BMI at the lower part of the distribution but a positive relationship with BMI at the upper part of the distribution. Indeed, in large datasets with sufficient numbers of thin participants, quantile regression estimates show reversal of the SEP-BMI association in the lower and upper end of the BMI distribution.^13^ Similarly, physical activity might reduce fat mass but increase muscle mass,^21 22^ such that physical activity might be related to lower body weight at the upper part of the distribution but related to higher body weight at the lower part of the distribution (due to the primary aim or effect of exercise being muscle gain/preservation rather than fat loss). A range of environmentally-attributable risk factors may have a stronger causal effects at the upper end of the BMI distribution—a recent twin study^23^ suggested that environmental effects on BMI may be stronger at the upper end, yet estimates of genetic effect stronger at the center of the distribution.

### 2. Risk factor confounding

Different effects sizes across the outcome distribution could be explained by the risk factor not measuring the construct of interest equivalently across the outcome distribution, or being confounded by other factors. For example, it is theoretically possible that individuals from low childhood social class backgrounds who have low BMI (rather than the anticipated high BMI), may in fact be a selected subset of participants who in fact are of higher SEP by some other measure (such as lower maternal education and/or family income). This would lead to spuriously weaker associations at lower quantiles matching those observed in Table 1, driven by low correlations between the SEP indicator used and the construct of interest. We recommend that researchers test this possibility, for example by examining the convergent validity of the exposure across the outcome distribution. In our data, we did so by examining associations between father’s social class at birth and maternal education across BMI quintiles—reassuringly, correlations were found across the BMI distribution and were in fact stronger at lower quantiles (Spearman’s R (from lowest-highest BMI quintiles=0.41, 0.39, 0.32, 0.35, 0.25)). Confounding by adult height is also a possibility, although an unlikely explanation for our findings; in linear regression models, height is not correlated with BMI, yet it is negatively associated at upper quantiles (Appendix Table 2).

### 3. Risk-factor modification

Risk factors for obesity tend to cluster and do not act in isolation.^24^ Unmeasured factors may modify the effect of the risk factor and lead to larger effects at the higher end of the distribution. For example, individuals with higher BMI values are more likely, than individuals with lower BMI values, to have genetic variants that cause excessive weight gain. As supported by the gene-by-environment literature, this genetic risk may result in the effect of poor diet, for example, being greater among individuals with higher BMI who have higher genetic risk for obesity. Environmental or behavioural factors could also work in the same way. For example, individuals with higher BMI values may live in areas with poorer dietary options such that the effects of socioeconomic disadvantage are more pronounced at the upper end of the distribution. Some findings appear to support this suggestion—for instance, in the UK Biobank, estimated effects of genes on BMI were larger amongst more deprived areas.^25^

### 4. Outcome scaling

Findings could be an artefact attributable to the scale of the outcome measure used. While it is possible that a given change in risk factor has a uniform effect across the BMI distribution—for example, a given diet intervention could lead to an equivalent 5kg/m^2^ loss for everyone exposed (ie, both those with average and high BMI values)—it may instead lead to a given percentage change (eg, 5%). When examined on the absolute (kg/m^2^) scale, a diet which uniformly affects percent change in weight would seem to have a larger effect at the upper end of the distribution, yet an identical effect when examined on the relative scale (5% of BMI=20= 1; 5% of BMI=30=1.5). Thus, it seems useful to examine the risk factor and BMI association at the upper end of the distribution on both absolute and relative scales. However, we demonstrate in Appendix Tables 2-3 that associations with the risk factors used and BMI are similar when BMI is modelled in relative (logged, %) terms.

## Implications and conclusions

Building on recent calls that researchers investigating descriptive trends in health examine both measures of average and distribution,^26^ we recommend that, in order to better understand the determinants of the distribution of population health, tools such as quantile regression could be used more frequently in aetiological epidemiology. This applies across many outcomes since population health (physical and mental) is ultimately thought to exist on a continuum.^2^ This is particularly so in obesity research, given the evidence for larger effect sizes at the upper parts of the BMI distribution and the limitations of conventional reliance on obesity cut-points, which leads to a loss of information and reduced statistical power. The uncertainty in the specific cut-points to use — particularly for direct measures of fat mass, and childhood BMI measures — is further motivation for its use.

How are understanding ‘distributional’ effects relevant for policy? If a risk factor has a causal effect on a health outcome, and its effect is heterogenous—with increasingly larger effects at higher values (where health is worse)—then intervening on this risk factor may have greater health benefits than anticipated than when only examined in average terms. Thus, this information may be useful to inform evidence-based policy decision-making, including on which interventions should be scaled up to promote health. Indeed, it has recently been suggested that clinical trials should report distributional changes in treatment groups in addition to reporting average differences.^27^ Alternatively, a risk factor may lead to lower average BMI due to differences in the lower part of the BMI distribution; given suggestions that BMI has a J-shaped relationship with mortality,^28^ such factors may have worse (or net negative) effects on population health than anticipated when considering average BMI values alone.

## Acknowledgments

We thank Dr Shaun Scholes and Dr Laura Howe for providing comments on an earlier version of this manuscript.

**Appendix Table 1.** Associations between risk factors for waist circumference (cm) at 45y using both linear regression and quantile regression in the 1958 British birth cohort study

| VARIABLES | Linear regression estimates<br>(mean difference in waist<br>circumference) | Quantile regression estimates<br>(difference in waist circumference at below quantiles) |  |  |  |  |
| --- | --- | --- | --- | --- | --- | --- |
|  |  | q10 | q25 | q50 | q75 | q90 |
| Paternal social class (per 1 category lower), birth | 0.45***<br>(0.14) | 0.29*<br>(0.17) | 0.50**<br>(0.22) | 0.22<br>(0.18) | 0.50**<br>(0.22) | 0.88***<br>(0.20) |
| Maternal weight (per 1 higher category), birth | 1.54***<br>(0.11) | 1.21***<br>(0.16) | 1.50***<br>(0.13) | 1.57***<br>(0.094) | 1.72***<br>(0.15) | 1.92***<br>(0.22) |
| General cognition (per 1 lower SDS), 11y | 1.65***<br>(0.18) | 1.35***<br>(0.23) | 1.61***<br>(0.30) | 1.72***<br>(0.21) | 1.73***<br>(0.24) | 2.15***<br>(0.38) |
| Physical exercise (inactive vs active), 42y | 1.62***<br>(0.37) | 0.63<br>(0.64) | 0.50<br>(0.56) | 1.43***<br>(0.50) | 2.47***<br>(0.37) | 2.53***<br>(0.73) |
| Observations | 7,016 | 7,016 | 7,016 | 7,016 | 7,016 | 7,016 |

Standard errors in parentheses
\*\*\* p<0.01, \*\* p<0.05, \* p<0.1
Note: models are mutually adjusted.

**Appendix Table 2.** Associations between risk factors for body mass index at 44/45y (logged) using both linear regression and quantile regression in the 1958 British birth cohort study, adjusted for adult height

| VARIABLES | Linear regression estimates<br>(mean difference in BMI) | Quantile regression estimates<br>(difference in BMI at below quantiles) |  |  |  |  |
| --- | --- | --- | --- | --- | --- | --- |
|  |  | q10 | q25 | q50 | q75 | q90 |
| Paternal social class (per 1 category lower), birth | 0.012***<br>(0.0017) | 0.010***<br>(0.0025) | 0.010***<br>(0.0025) | 0.011***<br>(0.0017) | 0.012***<br>(0.0027) | 0.020***<br>(0.0026) |
| Maternal weight (per 1 higher category), birth | 0.019***<br>(0.0014) | 0.016***<br>(0.0016) | 0.017***<br>(0.0011) | 0.017***<br>(0.0011) | 0.022***<br>(0.0017) | 0.024***<br>(0.0026) |
| General cognition (per 1 lower SDS), 11y | 0.014***<br>(0.0022) | 0.0081***<br>(0.0030) | 0.013***<br>(0.0028) | 0.016***<br>(0.0022) | 0.017***<br>(0.0034) | 0.020***<br>(0.0045) |
| Physical exercise (inactive vs active), 42y | 0.021***<br>(0.0047) | -0.0084<br>(0.0077) | 0.0056<br>(0.0052) | 0.021***<br>(0.0058) | 0.035***<br>(0.0051) | 0.039***<br>(0.0093) |
| Height at 45y (per 1m higher) | 0.029<br>(0.022) | 0.23***<br>(0.030) | 0.22***<br>(0.033) | 0.14***<br>(0.026) | -0.087***<br>(0.032) | -0.26***<br>(0.053) |
| Observations | 6,943 | 6,943 | 6,943 | 6,943 | 6,943 | 6,943 |

Standard errors in parentheses
\*\*\* p<0.01, \*\* p<0.05, \* p<0.1
Note: models are mutually adjusted.

**Appendix Table A3.**
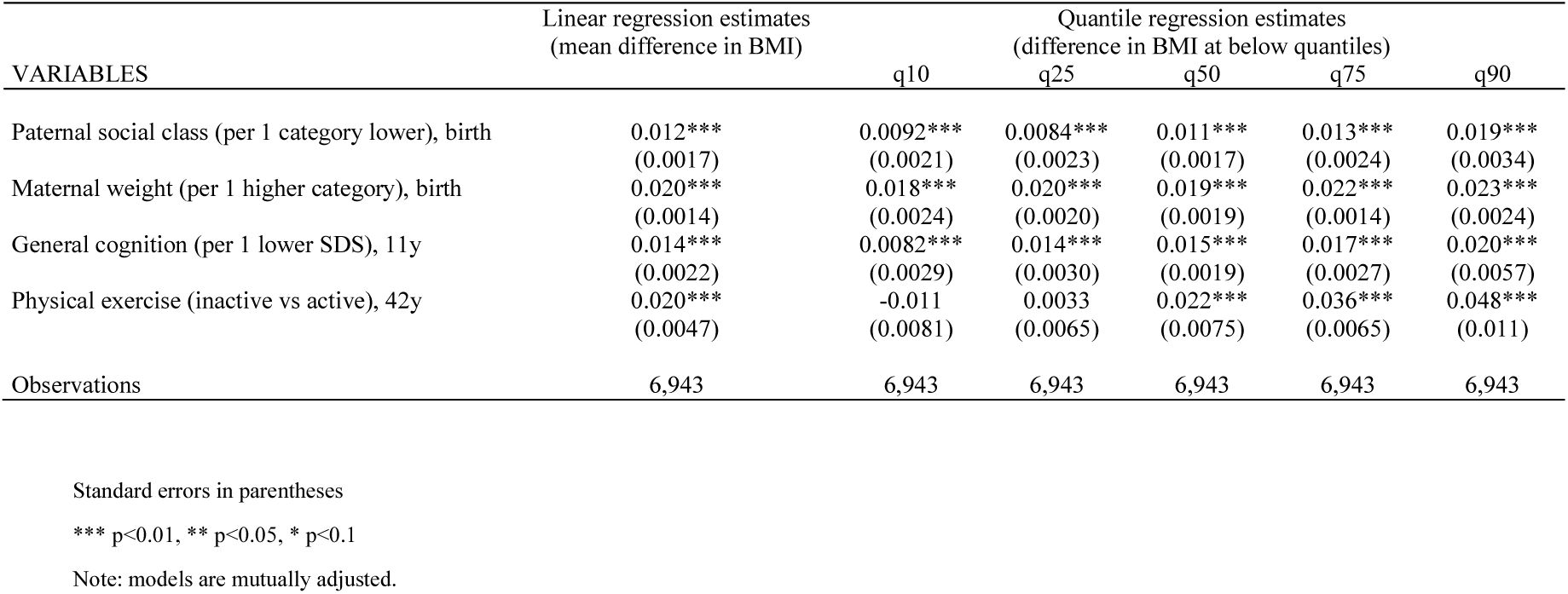
Associations between risk factors for body mass index at 45y (logged) using both linear regression and quantile regression in the 1958 British birth cohort study

